# Assessment of Neurovascular Uncoupling: *APOE* Status is a Key Driver of Early Metabolic and Vascular Dysfunction

**DOI:** 10.1101/2023.12.13.571584

**Authors:** Kristen Onos, Peter B. Lin, Ravi S. Pandey, Scott A. Persohn, Charles P. Burton, Ethan W. Miner, Kierra Eldridge, Jonathan Nyandu Kanyinda, Kate E. Foley, Gregory W. Carter, Gareth R. Howell, Paul R. Territo

## Abstract

**BACKGROUND:** Alzheimer’s disease (AD) is the most common cause of dementia worldwide, with apolipoprotein ε4 (*APOE****^ε^****^4^*) being the strongest genetic risk factor. Current clinical diagnostic imaging focuses on amyloid and tau; however, new methods are needed for earlier detection.

**METHODS:** PET imaging was used to assess metabolism-perfusion in both sexes of aging C57BL/6J, and h*APOE* mice, and were verified by transcriptomics, and immunopathology.

**RESULTS:** All h*APOE* strains showed AD phenotype progression by 8 mo, with females exhibiting the regional changes, which correlated with GO-term enrichments for glucose metabolism, perfusion, and immunity. Uncoupling analysis revealed *APOE****^ε^****^4/^****^ε^****^4^* exhibited significant Type-1 uncoupling (↓ glucose uptake, ↑ perfusion) at 8 and 12 mo, while *APOE****^ε^****^3/^****^ε^****^4^* demonstrated Type-2 uncoupling (↑ glucose uptake, ↓ perfusion), while immunopathology confirmed cell specific contributions.

**DISCUSSION:** This work highlights *APOE****^ε^****^4^* status in AD progression manifest as neurovascular uncoupling driven by immunological activation, and may serve as an early diagnostic biomarker.

## BACKGROUND

The incidence of Alzheimer’s disease (AD) and related dementias (RD) continues to rise globally [1]. Despite an increase in funding for research, and advancements in imaging technologies, which have largely focused on amyloid (Aβ) and neurofibrillary tau tangles (NFT) [2], it is still unclear what leads to neurodegeneration in the aging brain. Moreover, clinical diagnosis remains challenging, with few reliable biomarkers beyond Aβ and NFT, thus greater attention is needed in the development of a broader range of blood- and imaging-based readouts.

For imaging, the most commonly used positron emission tomography (PET) tracers detect amyloid (i.e. ^18^F-florbetapir, ^18^F-flutemetamol) or tau (i.e. ^18^F-flortaucipir). As more is learned about early risk factors for AD, the timing of these pathologies, and how they relate to cognition, changes in these tracers may come much too late to allow significant intervention [3]. Clinical studies have demonstrated a reduction in glucose brain metabolism and cerebral blood flow disturbances in at-risk patient populations even before detectable levels of amyloid accumulation [4–6]. This has led to the proposal of ^18^F-fluorodeoxyglucose (^18^F-FDG), a substrate for glucose uptake, to be added to the typical ATN diagnostic framework [2]. Clinical studies have demonstrated a reduction in ^18^F-FDG signal in multiple brain regions of AD patients, correlating with faster cognitive decline and brain atrophy [7]. Furthermore, patients with mild cognitive impairment (MCI) that also exhibit diminished ^18^F-FDG are more likely to progress to AD [8]. Currently there has been much debate regarding whether changes in ^18^F-FDG signal can be related to specific cell types [9]. Initially it was thought that these changes were driven primarily by neurons [10–13]; however, recent studies has shifted attention to glial cells, as they may contribute to the overall net ^18^F-FDG uptake [9]. Despite these advances, more work is necessary to fully appreciate cell-specific contributions that underlie ^18^F-FDG PET.

Under physiological stress, the brain may take different steps to cope, mitigate damage, or preserve homeostasis in response to an energy deficit [13]. This includes compensatory hyperemia, followed by a strong angiogenic response to counteract low oxygen and nutrients, thus leading to an increase in vascular density [14]. Arteriogenesis can also occur, driven by hemodynamic factors such as stretch and shear stress [15]. Disordered vascular remodeling and arteriovenous malformation can lead to blood vessel rupture and organ hemorrhage, and issues with brain perfusion have been identified in AD and other neurodegenerative disorders [16, 17]. Recent neuropathological studies have indicated that the majority of AD cases are comprised of mixed pathologies [18], with small vessel dysfunction being the most common [19].

Several genetic factors have been identified to confer AD risk; however, the ε4 allele on the *apolipoprotein E* (*APOE)* gene locus is the strongest for late-onset AD (LOAD). A single copy of the *APOE****^ε^****^4^* allele increases risk by 3-4 folds, while two copies augment the risk to 13 fold [20]. As study populations become more diverse, recent work has suggested that these odds ratios are limited to those of European descent. One of the many roles of APOE is regulating cerebrovascular integrity [21], with lack of APOE expression leading to blood-brain barrier (BBB) disruption [22, 23]. *APOE****^ε^****^4^* has been linked to decreased vascular density [24, 25], and is correlated with decreased cognitive performance. APOE has also been implicated in cerebral glucose metabolism [26], with ε4 identified as independent risk factors for T2D [27, 28] and cardiovascular disease [29, 30].

In this study, we sought to understand brain metabolism and perfusion changes at young and middle age time points in mice carrying combinations of humanized *APOE****^ε^****^3^* and *APOE****^ε^****^4^* alleles. We employed translational ^18^F-FDG for glucose uptake and ^64^Cu-PTSM for perfusion, and combined these into a novel measure of neurovascular coupling. We determined significant age, sex- and allele-specific differences across multiple regions in the brain. While imaging in the clinic is often cost-prohibitive and limited in rural settings, we believe that the ability to detect changes in neurovascular uncoupling could serve as an early diagnostic biomarker in at risk populations.

## METHODS AND MATERIALS

### Mouse Strains

The B6J.*APOE3* KI (h*APOE****^ε^****^3^*, available as B6.Cg-*Apoe^em2(APOE*)Adiuj^*/J, JAX#029018, the Jackson Laboratory (JAX)) and B6J.*APOE4* KI (h*APOE****^ε^****^4^*, available as B6(SJL)-*Apoe^tm1.1(APOE*4)Adiuj^*/J, JAX#027894, the Jackson Laboratory) mice strains created at the Jackson Laboratory carries a humanized APOE knock-in allele, in which a portion of the mouse Apoe gene (exons 2, 3, a majority of exon 4, and some 3’ UTR sequence) was replaced corresponding sequences of the human APOE isoform genes (**ε**3 and **ε**4). To generate h*APOE****^ε^****^3 /^****^ε^****^4^*, these two strains were intercrossed. Additional information on these mice are available from the Jackson Laboratory strain datasheets (h*APOE****^ε^****^3^*, https://www.jax.org/strain/029018; h*APOE****^ε^****^4^*, https://www.jax.org/strain/027894)

### Animal housing conditions

All experiments were approved by the Institutional Animal Care and Use Committee at Indiana University (IU) and at The Jackson Laboratory (JAX). Mice were bred in the mouse facility and maintained in a 12/12-hour light/dark cycle and room temperatures were maintained at 18-24°C (65-75°F) with 40-60% humidity. All mice were housed in positive, individually ventilated cages (PIV). Standard autoclaved 6% fat diet, (Purina Lab Diet 5K52) was available to the mice *ad lib*, as was water with acidity regulated from pH 2.5-3.0. All breeder and experimental mice were housed in the same mouse room and were aged together. Animals generated at JAX were aged and then shipped to IU for imaging within the week. All subjects were randomized and counterbalanced for testing order across multiples of instrumentation and time of day for each test day, with a simplified testing ID number (e.g., #1-100), with all technicians blinded to genotype (e.g., coded as A, B, C, etc.). The blind was maintained throughout testing and until after the data were analyzed with no subjects or data excluded based on any mathematical outliers.

### Magnetic Resonance Imaging (MRI)

To provide high contrast grey matter images, at least two days before PET imaging, mice were induced with 5% isoflurane (balance medical oxygen), placed on the head coil, and anesthesia maintained with 1-3% isoflurane for scan duration. High-resolution T2-weighted (T2W) MRI images were acquired using a 3T Siemens Prisma clinical MRI scanner outfitted with a dedicated 4 channel mouse head coil and bed system (RapidMR, Columbus OH). Images were acquired using a SPACE3D sequence [31] using the following acquisition parameters: TeA: 5.5min; TR: 2080ms; TE: 162ms; ETL: 57; FS: On; Ave: 2; Excitation Flip Angle: 150; Norm Filter: On; Restore Magnetization: On; Slice Thickness 0.2mm: Matrix: 171×192; FOV: 35×35mm, yielding 0.18 x 0.18 x 0.2mm resolution images. After the imaging period, mice were returned to their warmed home cages and allowed to recover.

### Radiopharmaceuticals and Study Population

Regional brain glycolytic metabolism was monitored using 2-[^18^F]-fluoro-2-deoxy-D-glucose (^18^F-FDG) and was synthesized, purified, and prepared according to established methods [32], where clinical unit doses ranging from 185 to 370 MBq (5 to 10 mCi) were purchased from PETNet Indiana (PETNET Solutions Inc). To evaluate region brain perfusion, Copper(II) pyruvaldehyde bis(N4-methylthiosemicarbazone) labeled with ^64^Cu (^64^Cu-PTSM) was synthesized, purified, and unit doses (i.e., 370 to 740 MBq (10 to 25 mCi)) dispensed by the PET Radiochemistry Core Facility at Washington University according to methods described previously [33, 34].

### Positron Emission Tomography (PET) and Computed Tomography (CT) Imaging

To evaluate changes in cerebral glycolysis (^18^F-FDG) and cerebral perfusion (^64^Cu-PTSM) mice were placed in a restrainer and consciously injected into the peritoneal or tail vein, respectively, with 3.7-11.1 MBq (0.1-0.3 mCi) of purified, sterile radiotracer, where the final volume did not exceed 10% of the animal’s body weight. Each animal was returned to its warmed home cage and allowed 30 min (^18^F-FDG) or 5 min (^64^Cu-PTSM) to allow for uptake and cellular trapping [35, 36]. Animals were fasted overnight only for imaging with ^18^F-FDG. Post-uptake, mice were induced with 5% isoflurane gas, placed on the scanner imaging bed, and anesthesia maintained at 1-3% isoflurane (balance medical oxygen) during acquisition. In all cases, calibrated PET acquisition was performed in list mode for 15 (^18^F-FDG) or 30 (^64^Cu-PTSM) min on an IndyPET3 scanner [37], where random prompts did not exceed 10% of the total prompt rate. Post-acquisition, the images were reconstructed into a single-static image with a minimum field of view of 60 mm using filtered-back-projection (FBP) and were corrected for decay, random coincidence events, and dead-time loss [38]. For some cohorts, both anatomical structure and function PET/CT imaging were performed with a Molecubes β-X-CUBE system (Molecubes NV, Gent Belgium), where calibrated list-mode PET images were reconstructed into a single-static image using ordered subset expectation maximization (OSEM) with 30 iterations and 3 subsets [39]. To provide anatomical reference and attenuation maps necessary to obtain fully corrected quantitative PET images, helical CT images were acquired with tube voltage of 50 kV, 100 mA, 100 μm slice thickness, 75 ms exposure, and 100 μm resolution. For β-CUBE studies, images were corrected for radionuclide decay, tissue attenuation, detector dead-time loss, and photon scatter according to the manufacturer’s methods [39].

### Image Processing and Analysis

All PET, CT and MRI images were co-registered using a ridged-body mutual information-based normalized entropy algorithm [40] with 9 degrees of freedom, and mapped to stereotactic mouse brain coordinates [41] using Analyze 12 (AnalyzeDirect, Stilwell KS) and MIM Encore Software 7.3.2 (Beachwood OH). Imaging Study data was collected and managed using RedCap electronic data capture tools hosted at Indiana University. Post-registration, 56 regions bilateral regions were extracted via brain atlas and left/right averaged to yield 27 unique volumes of interest that map to key cognitive and motor centers. To permit dose, scanner and brain uptake normalization, Standardized Uptake Value Ratios (SUVR) relative to the cerebellum were computed for PET for each subject, genotype, and age as follows:

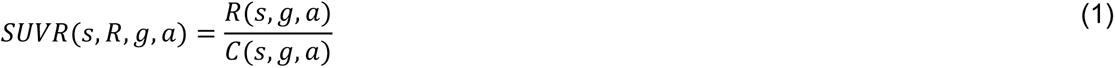

where, *s, g, a, R*, and *C* are the subject, genotype, age, region/volume of interest, cerebellum region/volume of interest. The SUVR values were then converted to z-score as follows:

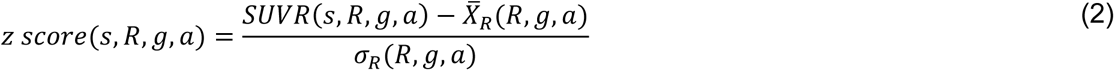

where, *s, g, a, R, X̄_R_*, and σ_*R*_ are the subject, genotype, age, mean of the reference population in SUVR, standard deviation of the reference population, based on the specified analytical strategies (effects of aging, humanized genes, and AD-risk alleles). Data are then projected onto Cartesian space, where the x-axis represents the z-score change in perfusion, derived from the ^64^Cu-PTSM data, and the y-axis is the z-score change in glucose uptake, via ^18^F-FDG, as a surrogate readout for glycolytic metabolism [12, 42, 43].

### Transcriptomic Analysis

RNA-Seq data (counts in transcript per kilobase million (TPM)) from cortex brain samples from four and 12 months old mice expressing hAPOE**^ε^**^3/^**^ε^**^3^ and hAPOE**^ε^**^4/^**^ε^**^4^ [44] were obtained from the AD Knowledge Portal (https://www.synapse.org/#!Synapse:syn26561824. We also obtained RNA-Seq data (counts in transcript per million) from whole left hemisphere brain samples from mice expressing human hAPOE**^ε^**^4/^**^ε^**^4^ [45] from the AD Knowledge Portal https://www.synapse.org/#!Synapse:syn17095983).

To start, we performed differential expression (DE) analysis using DESEq in R between hAPOE**^ε^**^4/^**^ε^**^4^ and B6 mice at 8 months. We calculated average Z-scores from imaging data across all 27 brain regions for ^18^F-FDG and ^64^Cu-PTSM. This was then followed by computing Pearson correlations between these average z-scores and DE for hAPOE**^ε^**^4/^**^ε^**^4^ and B6 mice. Next, we performed Gene Ontology (GO) term enrichment analysis on the genes positively correlated with ^18^F-FDG and ^64^Cu-PTSM. GO enrichment was performed using the function enrichGO from the clusterProfiler package [46]. The significance threshold level for all enrichment analyses was set to p ≤ 0.05 using Benjamini-Hochberg corrected p-values.

We then repeated these analysis steps for other group comparisons. To assess longitudinal changes with age in hAPOE**^ε^**^4/^**^ε^**^4^ mice, we performed DE comparing 12 months relative to 8 months, 12 months relative to 4 months and 8 months relative to 4 months for both sexes. This was followed by calculation of average z-scores from imaging data for longitudinal changes in ^18^F-FDG and ^64^Cu-PTSM measures in hAPOE**^ε^**^4/^**^ε^**^4^ mice for both sexes (male and female). We then performed Pearson correlations between the combined DE from aging and the z-scores from the imaging data, followed by GO enrichment analysis.

For our last set of comparisons, we looked at DE in hAPOE**^ε^**^4/^**^ε^**^4^ mice relative to hAPOE**^ε^**^3/^**^ε^**^3^ mice at 4 and 12 months for both sexes (male and female). This was followed by computation of correlation with imaging data z-score and GO enrichment analysis.

### Neuropathology

Additional mice were generated by crossing hAPOE3/3 and hAPOE4/4 and immunohistological assessments of blood vessels, microglia and astrocytes were performed in female animals. Briefly, animals were aged to 8 months and then transcardically perfused with PBS, and brains were fixed with 4% PFA overnight. This was followed by 15% sucrose and 30% sucrose on subsequent nights, and brains were then frozen and sectioned at 25 m on a microtome. Floating brain sections were stored in cryoprotectant at 4C. For staining, tissues were first rinsed in 1000 µl of PBS and then blocked with 5% Normal Goat Serum (NGS) and 1% PBT at 4C on a shaker for one hour. The tissues were then washed 2x for 20 minutes in PBS, followed by antibody staining, Iba1-Rabbit (1:200 Wako, #019-19741 PAK 6839), CD31 – Rat (1:300 BD Biosciences #550274 9259767) and GFAP-Chicken (1:300 Origine 87987979) at 4C on shaker for one night. Next, tissues were washed 3x in PBS for 10 minutes at RT on shaker and incubated with Goat secondaries (1:1000 Invitrogen #A11036 568; 1:1000 Life Technologies #A110006 488; 1:1000 Invitrogen Channel 633.) diluted with 1%PBT for 2 hours. This was followed by 3x washes in PBS for 10 minutes at RT on shaker. Lastly, DAPI staining was performed, tissues were rinsed in PBS and then mounted on glass slides for imaging.

For quantification, whole slice imaging was performed blind to genotype on 3 sections per animal at 20x on a Leica Dmi8 fluorescence wide-field microscope with LAS-X Navigator software. Individual channel images were converted to tagged image format (tif) and quantification was performed using ImageJ. Briefly, images were converted to 8 bits per pixel, and threshold applied via the triangle method using 20, and 255 as lower and upper cut-points. Post threshold, individual cells were detected across the entire image using Particle Analysis, with the following parameters: Size = [1,10]; Circularity = [0,1] for glia and [0, 250] for vessels; Exclude edges = True; Include holes = True; Show = Ellipses. Quantitation of object counts and percent of whole area were determined for 3 replicates per genotype.

High resolution imaging of representative cortex and hippocampus was also performed at 20x on this same set of tissue using a Leica SP8 confocal microscope. Image processing was also performed in ImageJ.

### Statistical Analysis

For standard PET image analysis, region data were plotted and analyzed by multiple unpaired T-test, and since only a single factor was being compared, Bonferroni corrections were not applied using GraphPad Prism. For uncoupling analysis, statistical analyses were performed within strain across age series. Student’s T-test was employed on data sets differing by a single variable (e.g., tracer, age, sex, genotype, and brain region), and significance level was considered at p < 0.05. To determine uncoupling, Student’s T-test was computed between ^64^Cu-PTSM and ^18^F-FDG for a single variable (e.g., age, sex, genotype, and brain region). To provide anatomical reference, significant brain regions for ^64^Cu-PTSM, ^18^F-FDG, and uncoupling were projected onto stereotactic mouse brain coordinates [41], and overlaid with anatomical T1W MRI derived from the Allen Common Coordinate Framework [47]. To show directionality of change for ^64^Cu-PTSM and ^18^F-FDG, p-values were assigned the sign of the z-scores from which they were derived, while p-values for uncoupled were unchanged. For immunopathology, data were analyzed by unpaired One-Way ANOVA with multiple comparisons, and plotted using GraphPad Prism.

### Availability of Data and Materials

All the datasets are available via the AD Knowledge Portal (https://adknowledgeportal.synapse.org). The AD Knowledge Portal is a platform for accessing data, analyses, and tools generated by the Accelerating Medicines Partnership (AMP-AD) Target Discovery Program and other National Institute on Aging (NIA)-supported programs to enable open-science practices and accelerate translational learning. The data, analyses, and tools are shared early in the research cycle without a publication embargo on a secondary use. Data is available for general research use according to the following requirements for data access and data attribution (https://adknowledgeportal.synapse.org/DataAccess/Instructions).

## RESULTS

### C57BL/6J Show Significant Sex- and Aging-Relevant Changes In Brain Metabolism and Perfusion

We first performed imaging on C57BL/6J (B6) mice in aged cross-sectional cohorts, as this is the genetic context in which the humanized APOE allelic series was introduced by MODEL-AD. For these analyses, we averaged the signal between left and right hemispheres, computed the standardized uptake value ratio (SUVR) based upon signal in cerebellum and assessed 27 major brain regions (Eqn. 1). Animals were fasted overnight and first imaged with ^18^F-FDG. Overall, both female and male B6 animals exhibited significant increases in ^18^F-FDG signal between 4 and 8 months in a similar set of regions (**Fig. 1A-B**): Dorsolateral Orbital cortex (DLO), Frontal Association cortex (FrA), Lateral Orbital cortex (LO), Medial Orbital cortex (MO), Prelimbic cortex (PrL), Primary motor cortex (M1), Secondary motor cortex (M2), and Ventral Orbital cortex (VO). Female B6 mice also exhibited significant increases between 4 and 8 months in Cingulate cortex (Cg), Corpus Callosum (CC), and Fornix (FN). Male B6 also exhibited significant increases between 4 and 8 months in Primary Somatosensory cortex (S1), and a significant decrease in the Dorso-Lateral-Intermedial-Ventral Entorhinal cortex (DLIVEnt). In comparisons performed between 8 and 12 months, female and male B6 mice demonstrated significant differences in ^18^F-FDG signal in independent regions. Females showed a decrease in Caudate Putamen (CPu), Cg, CC, DLO, FN, LO, MO and Thalamus (TH). Males exhibited an increase in DLIVEnt, but decreases in all regions of the Parietal cortices (PtPR and PtA), M1, Retrosplenial Dysgranular cortex (RSC), M2 and Primary and Secondary Visual cortex (V1V2). Finally, only female B6 displayed significant changes when comparing young with middle-aged animals (4 vs 12 months). In this comparison, a significant decrease was demonstrated in DLIVEnt, while there was a significant increase in S1 (see Supplemental Table 1).

**FIGURE 1:**
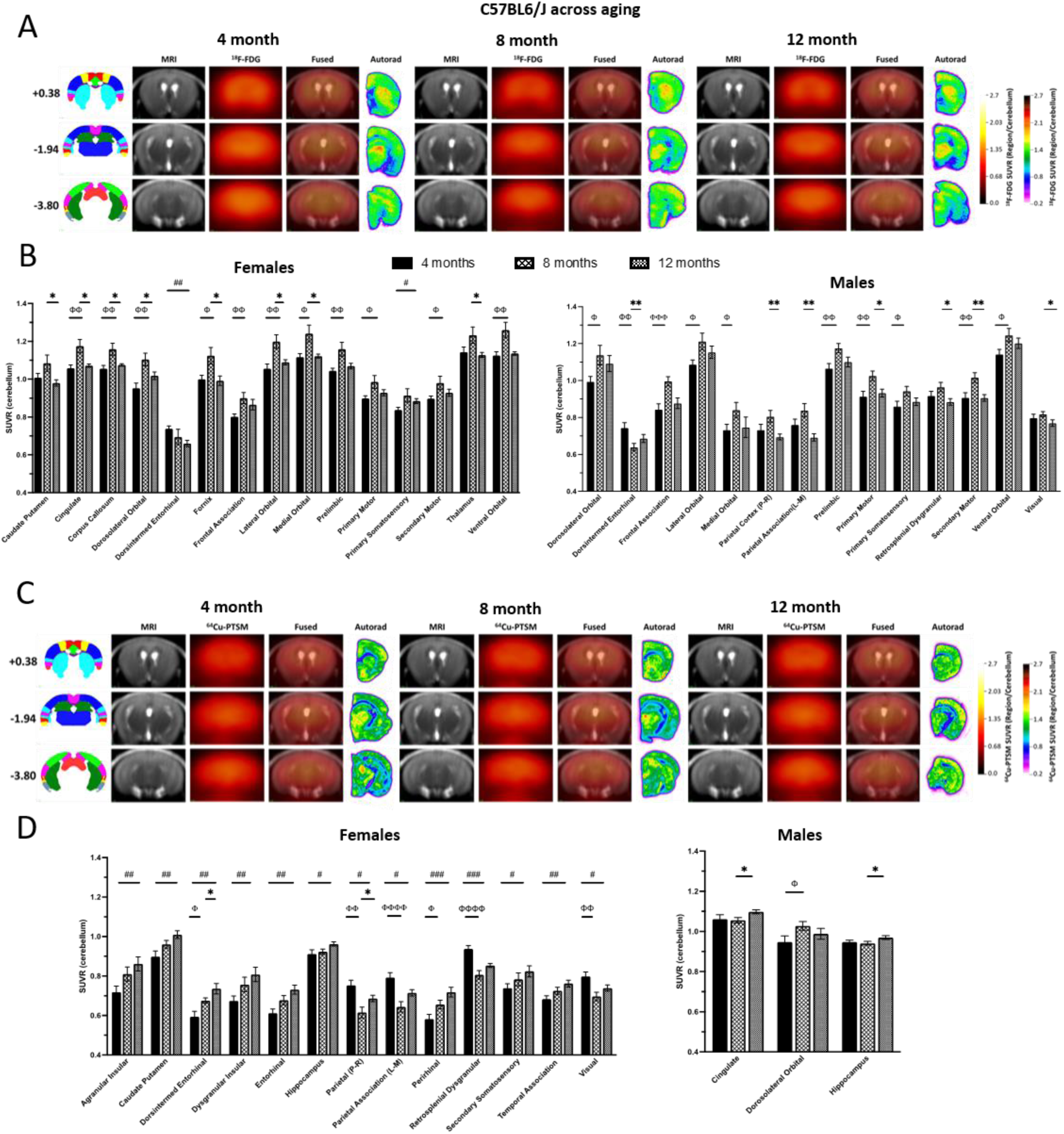
^18^F-FDG and ^64^Cu-PTSM in C57BL6/J across aging. (**A**) Representative MRI, 18F-FDG PET signal, fused PET signal and MRI, and AUTORAD at three bregma targets (Anterior +0.38, Medial -1.94 and posterior -3.80) for female C57BL6/J (B6) mice. (**B**) Computed standardized uptake value ratio (SUVR) relative to cerebellum is plotted in brain regions in male and female B6 mice that demonstrated a significant difference in ^18^F-FDG in one of the three age comparisons. Filled bars represent 4 months, lattice-patterned bars represent 8 months and checkered bars represent 12 months. Significant comparisons between 4 and 8-months are designated with Φ. Significant comparisons between 8 and 12-months are designated with *. Significant comparisons between 4 and 12 months are designated with a #. Number of symbols represents significance level, one is p ≤ 0.05, two is p ≤ 0.01, three is p ≤ 0.001 and four is p ≤ 0.0001. (**C**) Representative MRI, ^64^Cu-PTSM signal, fused PET signal and MRI, and AUTORAD at previously defined bregma targets in female B6 mice. (**D**) Computed standardized uptake value ratio (SUVR) relative to cerebellum is plotted in brain regions in male and female B6 mice that demonstrated a significant difference in ^64^Cu-PTSM in one of the three age comparisons. Bar patterns, symbols and significance values are the same as B.

Imaging with ^64^Cu-PTSM occurred in the same animals after a recovery period to allow tracer decay and animal recovery from fasting. For ^64^Cu-PTSM, female B6 exhibited significant changes in many more brain regions than male B6 mice (**Fig. 1C-D**). Female mice demonstrated significant differences in perfusion between 4 and 8 months in DLIVEnt, PtPR, PtA, Perirhinal cortex (PRH), RSC and V1V2. Two regions continued to show significant changes between 8 and 12 months, DLIVEnt and PtPR. Ultimately, the greatest differences in the most regions occurred in comparisons between 4 and 12 months in Agranular Insular cortex (AI), CPu, DLIVEnt, Dysgranular Insular cortex (DI), Ectorhinal cortex (ECT), Hippocampus (HIPP), PtPR, PtA, PRH, RSC, Secondary Somatosensory cortex (S2), Temporal Association cortex (TeA) and V1V2. In male B6, regional changes were limited to an increase between 4 and 8 months in DLO, and increases between 8 and 12 months in Cg and HIPP.

### Metabolism and Perfusion Changes are Predominate in Female, Rather Than Male hAPOE^ε3/ε3^ mice

*APOE****^ε^****^3/^****^ε^****^3^* has been designated as the most common isoform combination in the human population and is typically used as the control or ‘WT’ genotype in studies assessing the role of APOE in AD risk. In line with this, we assessed male and female mice that were homozygous for h*APOE****^ε^****^3/^****^ε^****^3^*. Overall, sex was a significant driver of age-related changes, as female h*APOE****^ε^****^3/^****^ε^****^3^* carriers demonstrated more regions changed with ^18^F-FDG and PTSM PET in comparison to male mice (**Fig. 2**). In comparisons between 4 and 8 months for ^18^F-FDG, females exhibited an increase in FDG signal in the RSC and TH (**Fig. 2A-B**). In comparisons between 8 and 12 months, there was a significant reduction in FDG signal in the Auditory cortex (AuDMV) and Cg. A larger set of regions exhibited significant changes in ^18^F-FDG between 4 and 12 months; AI, AuDMV, CPu, Cg, DI, and S2 were decreased with age, while PtA and V1V2 were increased. In male h*APOE****^ε^****^3/^****^ε^****^3^* mice, CC, M1 and RSC increased ^18^F-FDG signal at 8 months relative to 4 months (**Fig. 2C-D**). There were no significant changes in ^18^F-FDG that occurred from 8 to 12 months, however, RSC and V1V2 did demonstrate significant increases in comparisons of 4 and 12 months.

**FIGURE 2:**
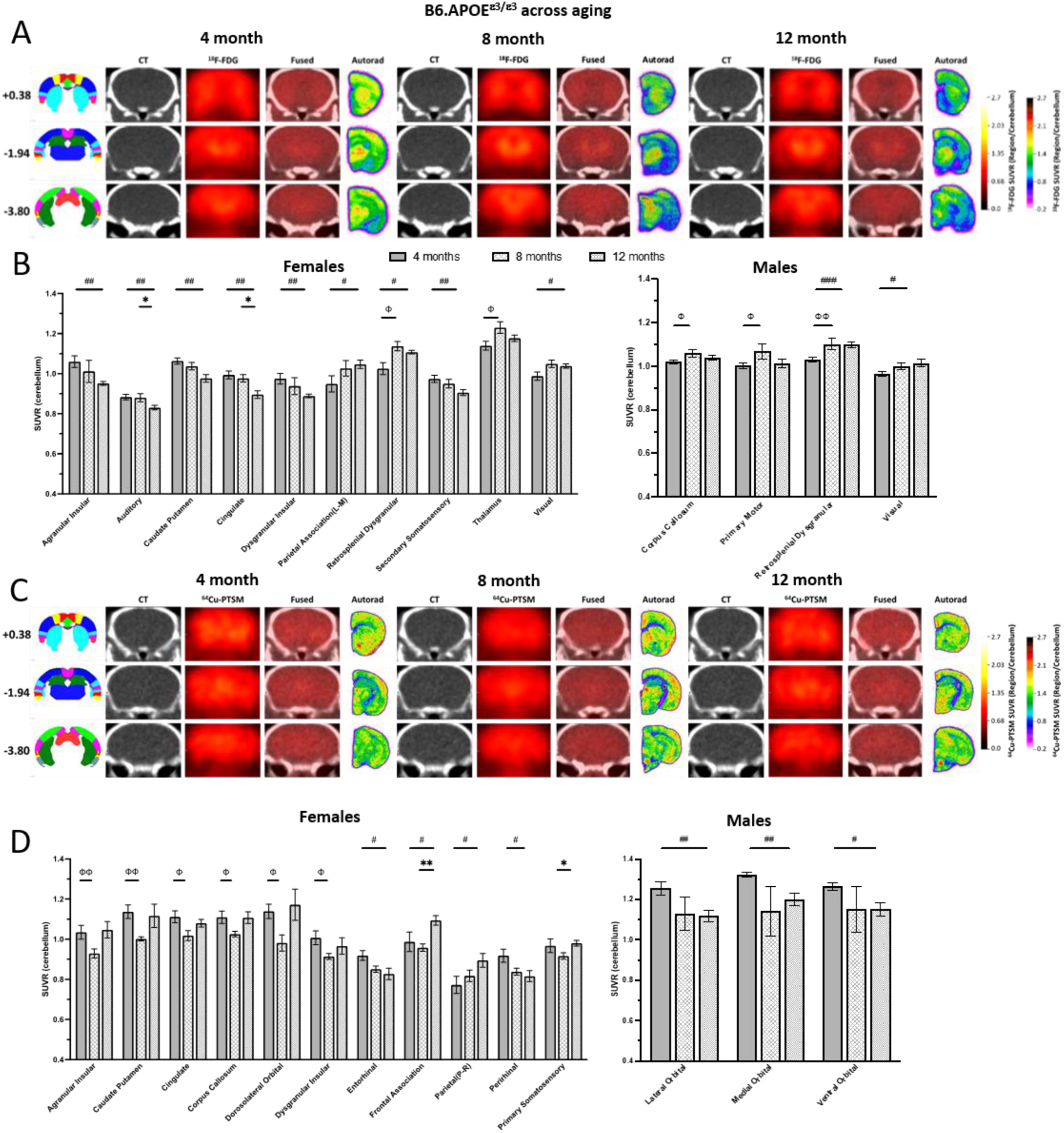
^18^F-FDG and ^64^Cu-PTSM in humanized APOE^ε3/ε3^ across aging. (**A**) Representative MRI, ^18^F-FDG PET signal, fused PET signal and CT, and AUTORAD at three bregma targets (Anterior +0.38, Medial -1.94 and posterior -3.80) for female humanized *APOE****^ε^****^3/^****^ε3^*** (h*APOE****^ε^****^3/^****^ε^****^3^*) mice. (**B**) Computed standardized uptake value ratio (SUVR) relative to cerebellum is plotted in brain regions in male and female h*APOE****^ε^****^3/^****^ε^****^3^* mice that demonstrated a significant difference in ^18^F-FDG in one of the three age comparisons. Filled bars represent 4 months, lattice-patterned bars represent 8 months and checkered bars represent 12 months. Significant comparisons between 4 and 8-months are designated with Φ. Significant comparisons between 8 and 12-months are designated with *. Significant comparisons between 4 and 12 months are designated with a #. Number of symbols represents significance level, one is *p* ≤ 0.05, two is *p* ≤ 0.01, three is *p* ≤ 0.001 and four is *p* ≤ 0.0001. (**C**) Representative CT, ^64^Cu-PTSM signal, fused PET signal and MRI, and Autorad at previously defined bregma targets in female h*APOE****^ε^****^3/^****^ε^****^3^* mice. (**D**) Computed standardized uptake value ratio (SUVR) relative to cerebellum is plotted in brain regions in male and female h*APOE****^ε^****^3/^****^ε^****^3^* mice that demonstrated a significant difference in ^64^Cu-PTSM in one of the three age comparisons. Bar patterns, symbols and significance values are the same as B.

For ^64^Cu-PTSM, female h*APOE****^ε^****^3/^****^ε^****^3^* mice demonstrated significant decreases in perfusion from 4 to 8 months in AI, CPu, Cg, CC, DLO and DI. In comparisons performed between 8 and 12 months, significant increases in ^64^Cu-PTSM were observed in FrA and S1. Finally, in comparisons between 4 and 12 months, ECT and PRH signals were decreased, while FrA and PtPR signals were increased. Male h*APOE****^ε^****^3/^****^ε^****^3^* carriers only exhibited significant changes in PTSM in three regions, LO, MO and VO, and only in comparisons between 4 and 12 months.

### Female hAPOE^ε4/ε4^ Mice Exhibit Significant Decreases in Metabolism and Perfusion in Almost Every Brain Region with Age

*APOE****^ε^****^4/^****^ε^****^4^* is predicted to exert effects in a multitude of ways that could compromise brain health, especially in aging. Regardless of sex, mice carrying h*APOE****^ε^****^4/^****^ε^****^4^* exhibited significant changes in ^18^F-FDG signal across the majority of brain regions tested, particularly in older ages (i.e. 12 months). In both B6 and h*APOE****^ε^****^3/^****^ε^****^3^*, brain regions showed significant increases in ^18^F-FDG signal between 4 and 8 months (**Fig. 1-2**). In contrast, both female and male h*APOE****^ε^****^4/^****^ε^****^4^* mice demonstrated significant decreases in most brain regions during this same time period (**Fig. 3A-B**). In female h*APOE****^ε^****^4/^****^ε^****^4^*, significant decreases in ^18^F-FDG were present in AuDMV, DLIVEnt, ECT, PtPR, PtA, PRH, RSC, S2, TeA and V1V2 between 4 and 8 months. In male h*APOE****^ε^****^4/^****^ε^****^4^*, these decreases occurred in AI, PtPR, PtA, RSC and V1V2, while the FrA exhibited a significant increase. In comparison between 8 and 12 months, females demonstrated increases (or approximate return to younger 4 month levels) in AI, AuDMV, DLIVEnt, DI, ECT, PRH, S2 and TeA. For male h*APOE****^ε^****^4/^****^ε^****^4^* carriers, this increase in FDG signal often surpassed levels previously seen at 4 months; AI, AuDMV, DLO, DLIVEnt, DI, ECT, FrA, HIPP, LO, PtPR, PRH, S1, S2, TeA and V1V2 all demonstrated significant increases in ^18^F-FDG signal at 12 months relative to 8 months. Lastly, in comparisons between 4 and 12 months, females showed significant decreases in ^18^F-FDG signal in Cg, CC, PtPR, PtA, RSC and V1V2. In contrast, males showed significant changes in ^18^F-FDG uptake between 4 and 12 months; AI, AuDMV, DLO, DLIVEnt, DI, ECT, FrA, PRH, S2 and TeA were all increased, while Cg and RSC were decreased.

**FIGURE 3:**
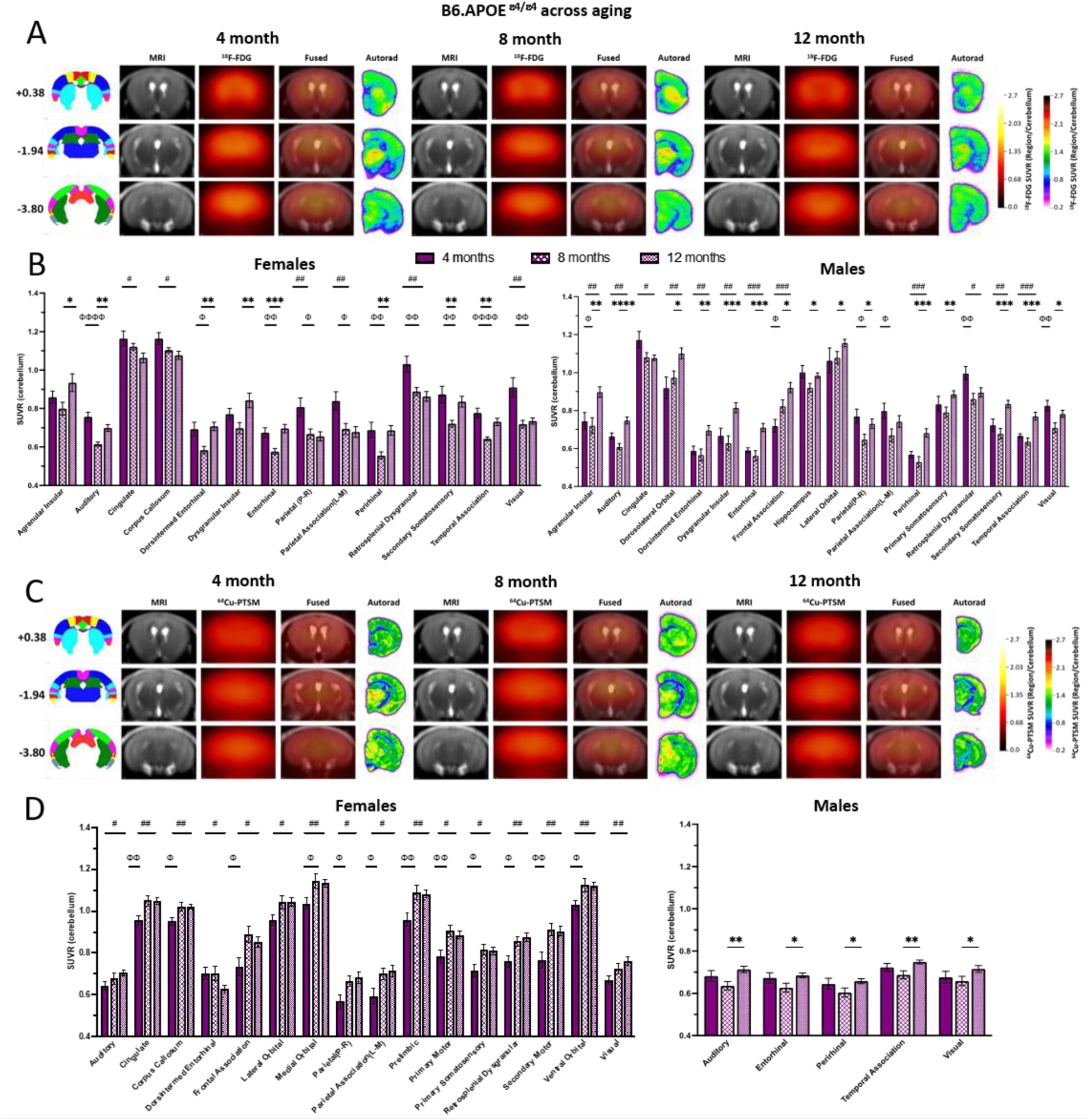
^18^F-FDG and ^64^Cu-PTSM in humanized APOE^ε4/ε4^ across aging. (**A**) Representative MRI, ^18^F-FDG PET signal, fused PET signal and MRI, and Autorad at three bregma targets (Anterior +0.38, Medial -1.94 and posterior -3.80) for female humanized *APOE****^ε^****^4/^****^ε^****^4^* (h*APOE****^ε^****^4/^****^ε^****^4^*) mice. (**B**) Computed standardized uptake value ratio (SUVR) relative to cerebellum is plotted in brain regions in male and female h*APOE****^ε^****^4/^****^ε^****^4^* mice that demonstrated a significant difference in ^18^F-FDG in one of the three age comparisons. Filled bars represent 4 months, lattice-patterned bars represent 8 months and checkered bars represent 12 months. Significant comparisons between 4 and 8-months are designated with Φ. Significant comparisons between 8 and 12-months are designated with *. Significant comparisons between 4 and 12 months are designated with a #. Number of symbols represents significance level, one is *p* ≤ 0.05, two is *p* ≤ 0.01, three is *p* ≤ 0.001 and four is *p* ≤ 0.0001. (**C**) Representative MRI, ^64^Cu-PTSM signal, fused PET signal and MRI, and AUTORAD at previously defined bregma targets in female h*APOE****^ε^****^4/^****^ε^****^4^* mice. (**D**) Computed standardized uptake value ratio (SUVR) relative to cerebellum is plotted in brain regions in male and female h*APOE****^ε^****^3/^****^ε^****^3^* mice that demonstrated a significant difference in ^64^Cu-PTSM in one of the three age comparisons. Bar patterns, symbols and significance values are the same as B.

Imaging with ^64^Cu-PTSM revealed significant increased signal in female h*APOE****^ε^****^4/^****^ε^****^4^* carriers between 4 months and 8 months, and this elevation often persisted into 12 months. This occurred in Cg, CC, FrA, MO, PtPR, PtA, PrL, M1, S1, RSC, M2, and VO. Many of the same regions were also significant for comparisons between 4 and 12 months, Cg, CC, FrA, MO, PtPR, PtA, PrL, M1, S1, RSC, M2, and VO. Additional regions in which significant increases in ^64^Cu-PTSM signal were determined were the AuDMV, DLIVEnt, LO and V1V2. In contrast, male h*APOE****^ε^****^4/^****^ε^****^4^* carriers demonstrated significant increases in ^64^Cu-PTSM only in comparisons between 8 and 12 months; these regions were AuDMV, ECT, PRH, TeA and V1V2.

### Correlation of Transcriptional Profiling with PET Tracer Outcomes Can Identify Critical Disease Processes

To elucidate the relationship between glycolytic metabolism and tissue perfusion with gene expression profiles, we ultilized two transcriptional profiling datasets. The first dataset consisted of hemi-coronal brains from 4, 8 and 12 month h*APOE****^ε^****^4/^****^ε^****^4^* and B6 animals that had been imaged in this study. The second data set was generated from dissected cortex and hippocampus from 4 and 12 month h*APOE****^ε^****^4/^****^ε^****^4^* and h*APOE****^ε^****^3/^****^ε^****^3^* animals. As the sample handling, library prep and sequencing were all performed at JAX for both datasets, we were interested in determining if there were differences in detection of transcripts (transcripts per kilobase million, TPMs) in h*APOE****^ε^****^4/^****^ε^****^4^* dependent on whether samples were hemi brain or individual brain regions (**Fig. 4A**). Overall, the detection of transcripts was similar in both female and male h*APOE****^ε^****^4/^****^ε^****^4^* mice regardless of tissue origin or age point, with a correlation coefficient of R = 0.97 comparing hemi brains to cortex or hippocampus (**Fig. 4B**). To further understand the overlap between these two datasets, we also performed correlations on the differentially expressed sets (DE; log2 fold change) produced by comparing 12 to 4 month h*APOE****^ε^****^4/^****^ε^****^4^* within each tissue (**Fig. 4C**). For these comparisons, the strongest correlations were between tissue type for each sex, rather than hemi-brains. For example, DE from cortices between females and males, and DE from hemi brains between females and males demonstrated the highest correlation, and there was no correlation between DE from hemi brains and cortex. Given this, we performed correlations between imaging and DE only within each of these datasets, rather than across. This meant that DE between B6 and h*APOE****^ε^****^3/^****^ε^****^3^* could not be compared to each other.

**FIGURE 4:**
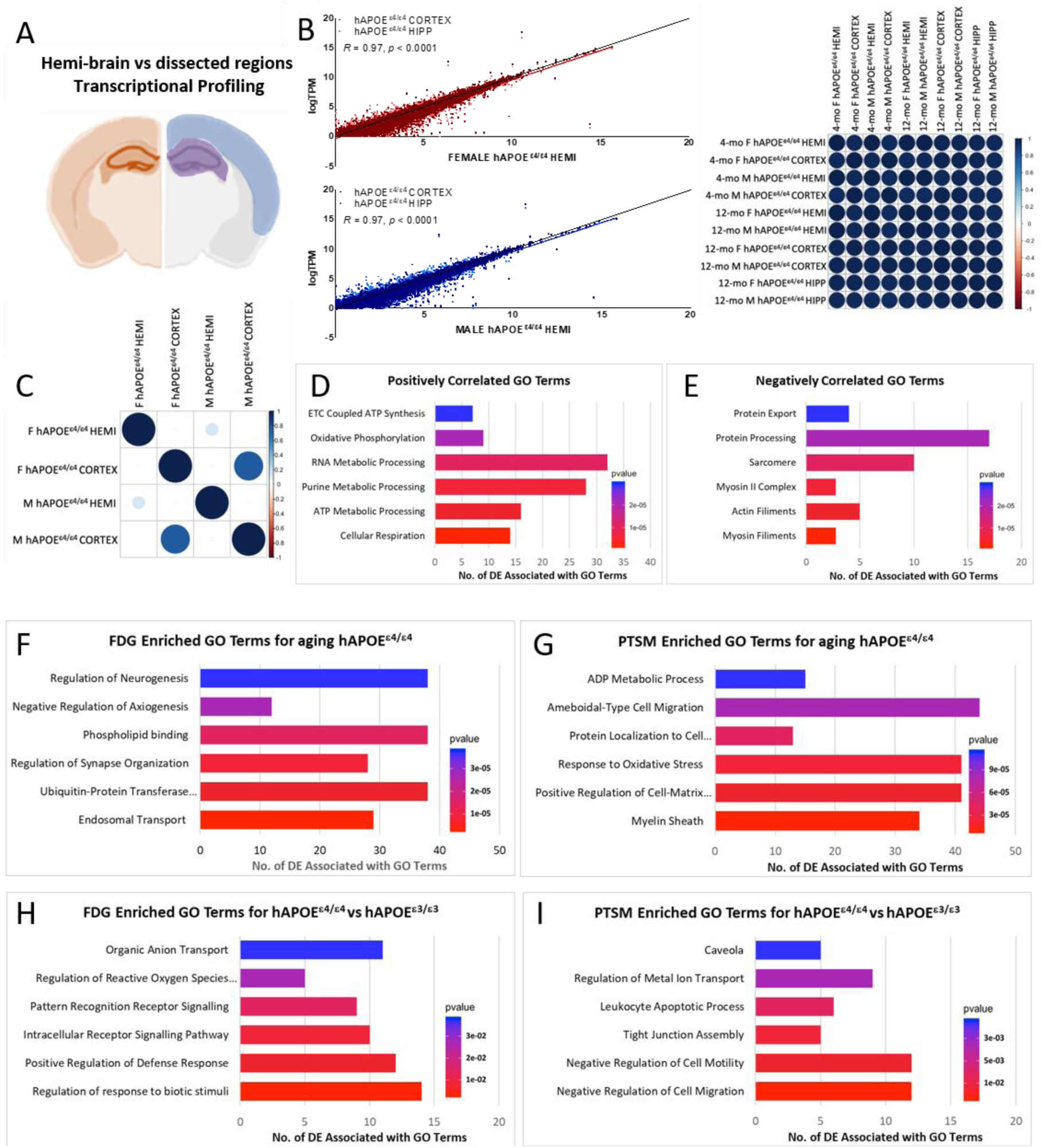
Correlation of differential gene expression with average whole brain ^18^F-FDG and ^64^Cu-PTSM signal. (**A**) Mouse brain where rust designates brain hemisphere (HEMI), blue designates dissected cortex and purple designates dissected hippocampus (HIPP) collected for two transcriptional profiling projects including aged h*APOE* mice. (**B**) Graphs that demonstrate the correlation between detected transcript expression from sequenced HEMI brains, cortex (dark red or dark blue) and HIPP (light red or light blue) in 12-month h*APOE****^ε^****^4/^****^ε^****^4^* females (top middle panel) and males (bottom middle panel). Correlation matrix (right) is also provided to demonstrate similarly across ages and regions in transcript expression data from the two sequencing projects. (**C**) Correlation matrix demonstrating the relationship between brain regions using differential expression of h*APOE****^ε^****^4/^****^ε4^*** between 12 and 4 months of age. (**D**) Comparing B6 and h*APOE****^ε^****^4/^****^ε4^*** averaged ^18^F-FDG PET signal revealed an enrichment of differentially expressed (DE) transcripts that were positively correlated with Gene Ontology (GO) terms relevant to cellular metabolism. ETC=Electron Transport Chain. (**E**) Comparing B6 and h*APOE****^ε^****^4/^****^ε4^*** averaged ^64^Cu-PTSM PET signal revealed DE enrichment that was negatively correlated with GO terms relevant to smooth muscle function, protein processing and export. (**F**) Comparing longitudinal age-related h*APOE****^ε^****^4/^****^ε4^*** averaged ^18^F-FDG PET signal identified an enrichment of differentially expressed (DE) transcripts that were positively correlated with GO terms relevant to neuronal function, lipid handling and targeted degradation. (**G**) Comparing longitudinal age-related h*APOE****^ε^****^4/^****^ε^****^4^* averaged ^64^Cu-PTSM PET signal identified an enrichment of differentially expressed (DE) transcripts that were correlated with GO terms relevant to cellular structure, migration and energy production. Full GO pathway name: Positive Regulation of Cell-Matrix Adhesion. (**H**) Comparing h*APOE****^ε^****^3/^****^ε^****^3^* and h*APOE****^ε^****^4/^****^ε4^*** averaged ^18^F-FDG PET signal determined an enrichment of DE transcripts that were correlated with GO terms relevant to signaling between cells in response to stimuli. (**I**) Comparing h*APOE****^ε^****^4/^****^ε4^*** and h*APOE****^ε^****^3/^****^ε^****^3^* averaged ^64^Cu-PTSM PET signal determined an enrichment of DE transcripts that were correlated with GO terms relevant to cellular movement, the blood brain barrier and immune-regulated cell death.

In order to compare gene expression and ^18^F-FDG or ^64^Cu-PTSM imaging, we averaged signal change between selected groups from all 27 brain regions, and then correlated this to DE. For example, comparisons performed between 8 month h*APOE****^ε^****^4/^****^ε^****^4^* and B6 revealed 587 transcripts that were significantly positively correlated (r>0.80, p<0.05) with ^18^F-FDG signal change, and 810 transcripts that were significantly negatively correlated (r<-0.80, p<0.05). When we performed Gene ontology (GO) term analysis, there was a significant enrichment of glycolytic and ox-phos pathways associated with cellular respiration (**Fig. 4D**). Similarly, analysis performed with ^64^Cu-PTSM signal change showed 346 transcripts that were significantly positively correlated (r>0.80, p<0.05) and 246 transcripts that were negatively correlated with perfusion measures (r<-0.80, p<0.05). GO term analysis showed there was a significant enrichment of protein export and processing, sarcomere and myosin filaments in negatively correlated transcripts (**Fig. 4E**).

Next, we were interested in longitudinal DE in h*APOE****^ε^****^4/^****^ε^****^4^* mice. For this analysis, we performed comparisons in 12 months relative to 8 months, 12 months relative to 4 months and 8 months relative to 4 months for both sexes. Then, we calculated average z-scores from imaging data across all 27 brain regions for longitudinal changes in ^18^F-FDG and ^64^Cu-PTSM measures in the same mice. This was followed by computing Pearson correlations between average z-scores for ^18^F-FDG and ^64^Cu-PTSM and longitudinal DE in h*APOE****^ε^****^4/^****^ε^****^4^* mice. We then performed GO term analysis which showed enrichment in pathways specific to neuronal function, lipid handling and targeted degradation for ^18^F-FDG (**Fig. 4F**), and enrichment in pathways specific to cellular structure, migration and energy production for ^64^Cu-PTSM (**Fig. 4G**).

Finally, we wanted to explore differences between h*APOE****^ε^****^3/^****^ε^****^3^* and h*APOE****^ε^****^4/^****^ε^****^4^* mice. Due to dataset limitations, this comparison could only be performed cross-sectionally between 4 and 12 months. Data analysis was performed the same as above. GO term analysis highlighted pathways specific to signaling between cells in response to stimuli for ^18^F-FDG (**Fig. 4H**), and pathways relevant to cellular movement, blood brain barrier integrity and immune-regulated cell death for ^64^Cu-PTSM (**Fig. 4I**).

### Combining ^18^F-FDG and ^64^Cu-PTSM PET Outcomes Into Early Disease Biomarkers of Neurovascular Uncoupling

To more completely understand the regional changes in perfusion and glycolytic metabolism, which make up neurovascular coupling, we performed bi-directional z-score analysis. This analysis (i.e. uncoupling analysis) was based on Eqn. 2, where z-scores for ^18^F-FDG and ^64^Cu-PTSM are projected onto Cartesian space with z-score changes in perfusion from the reference population plotted on the x-axis, and z-score changes in glycolysis from the reference population plotted on the y-axis, respectively. This approach establishes four primary quadrants (**Fig. 5A, Left**), with two of these showing coupled increases (Quadrant 1) or decreases (Quadrant 3) in perfusion and metabolism. These regions represent the normal physiological responses described as “coupled responses” for the neurovascular unit [13, 48, 49]. By contrast, the remaining two quadrants show uncoupled changes in perfusion and metabolism, with Quadrant 4, exhibiting decreases in metabolism, with increased hyperemia, and was referred to as Type 1 Uncoupling. Similarly, Quadrant 2 presents with an increase in metabolism, while perfusion is decreased relative to the reference population, and is referred to as Type 2 Uncoupling (**Fig. 5A, Left**). As the underlying data are plotted as z-scores, the changes along each axis represents the number of standard deviation changes away from the reference population, with a statistical probability of these distributions being different, such that ±1 = %68, ±2 = 95%, and ±3 = 99% (**Fig. 5A, Right**).

**FIGURE 5:**
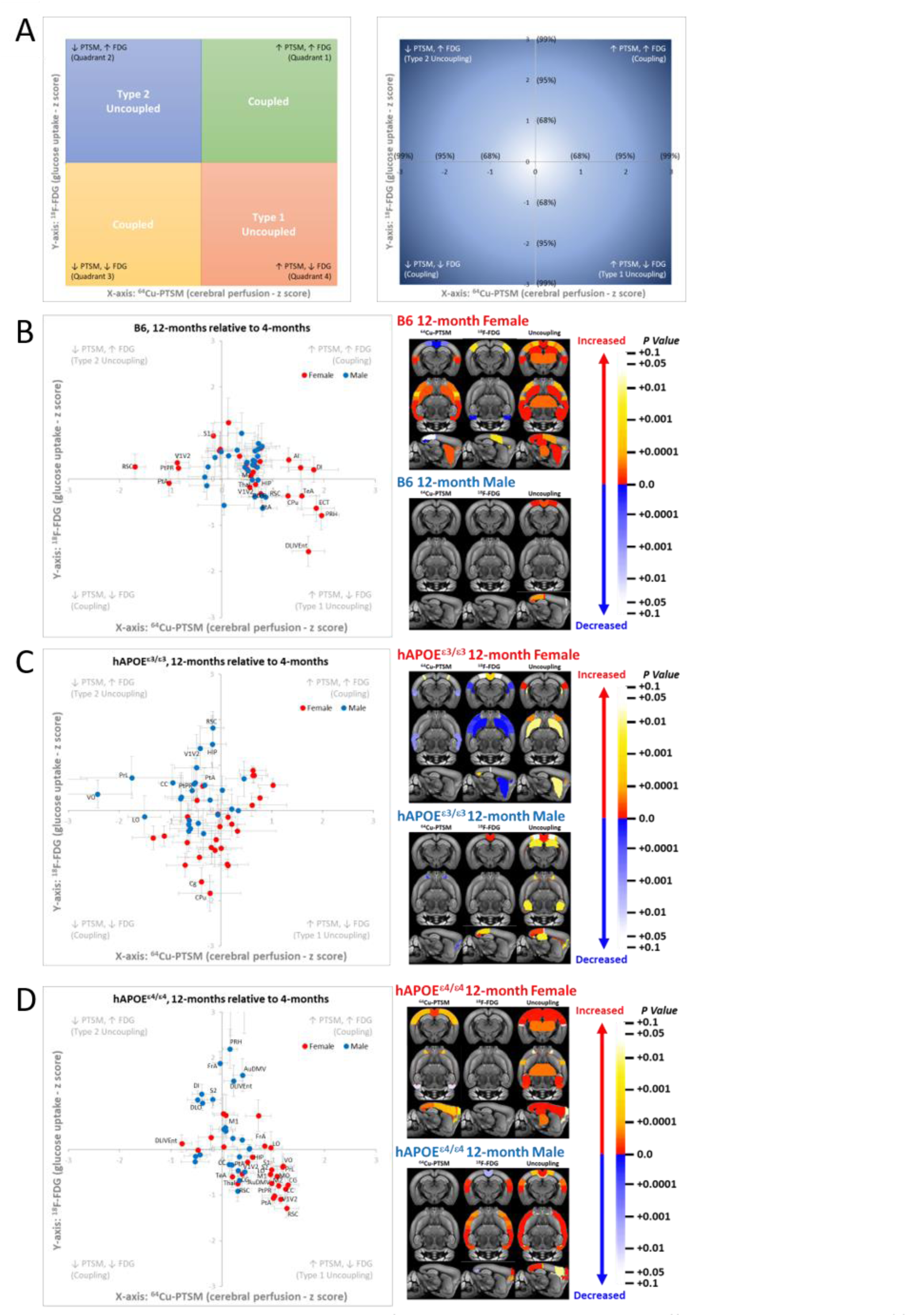
Assessment of Neurovascular Coupling at 12-months. (**A**) To enable assessment of neurovascular coupling (or uncoupling), comparisons can be made across appropriately selected groups by converting the ^18^F-FDG PET signal and ^64^Cu-PTSM PET signal to a z-score for each brain region, and then plotting these regions on the Y (^18^F-FDG) and X (^64^Cu-PTSM) axes. Brain regions that fall towards the center of this plot and within one standard deviation are likely functioning at a normal level. A brain region is determined to be coupled if both signals are in the same direction (increased or decreased), and uncoupled if they are opposite. (**B**) Analysis of neurovascular coupling of 12-month B6 in comparison with 4-month B6. Individual points represent the averaged z-scores of one of 27 brain regions, with red designating female and blue designating male. Image panels denote brain regions in the coronal, ventral and sagittal planes that exhibit significant differences in with each tracer, or the combined z-scores for uncoupling. Warmer colored regions indicated an increase, while cooler colored regions indicate a decrease. (**C**) of 12-month _h*APOE*_***ε****3/****ε****3* in comparison with 4-month h*APOE****^ε^****^3/^****^ε^****^3^*. (**D**) Analysis of neurovascular coupling of 12-month h*APOE****^ε^****^4/^****^ε^****^4^* in comparison with 4-month h*APOE****^ε^****^4/^****^ε^****^4^*. Female h*APOE****^ε^****^4/^****^ε^****^4^* exhibit a significant number of regions that are Type 1 uncoupled.

To maximize the likelihood of observing a change in z-score across each genotype, we performed longitudinal uncoupling analysis on 12 month mice, where each cohort referenced 4 month mice, which are sexually mature and without disease. As a baseline, we explored how mouse APOE in B6 mice affects perfusion and metabolism with age. Consistent with our overarching hypothesis, B6 mice at 12 months showed alterations in perfusion and metabolism which were sexually dimorphic (**Fig. 5B**), with male mice showing no statistically significant changes in ^64^Cu-PTSM or ^18^F-FDG, while the following regions: M2, PtA, PrL, PtPR, RSC, and V1V2 were uncoupled. In contrast, female B6 mice showed a much larger number of regions (i.e. AI, CPu, DLIVEnt, DI, ECT, HIP, PtPR, PtA, PRH, S1, RSC, TeA, TH, V1V2) (14/27) that exhibited significant changes in both tissue perfusion and Type 2 uncoupling (decreased perfusion, increased glucose uptake) (**Fig. 5B**).

To understand the impact of h*APOE****^ε^****^3/^****^ε^****^3^* on neurovascular coupling, both male and female mice were subjected to uncoupling analysis at 12 months. Unlike the B6 cohort, the addition of h*APOE****^ε^****^3/^****^ε^****^3^* resulted in minimal changes in perfusion or metabolism changes for both sexes (**Fig. 5C**). The number and degree of significantly uncoupled regions was greater for males than females at this age. Interestingly, male mice showed significant uncoupling in: CC, HIP, LO, MO, PtPR, PtA, PrL, RSC, VO, and V1V2, while female mice showed significant uncoupling in the AI, AuDMV, CPu, and Cg cortices.

To explore the disease-associated isoform of APOE, which is known to increase the likelihood of both vascular and metabolic dysfunction with age, mice bearing h*APOE****^ε^****^4/^****^ε^****^4^* at 12 months were analyzed for neurovascular uncoupling. Similar to non-disease isoform of APOE, male mice showed significant changes primarily in regional metabolism at this age, while female mice showed a significant increase in tissue perfusion (**Fig. 5D**). This sexual dimorphism in physiological changes in perfusion and metabolism resulted in a very large number of brain regions in both male (i.e. AuDMV, Cg, CC, DLO, DLIVEnt, DI, ECT, FrA, PtA, PRH, M1, RSC, S2, TeA, and V1V2) and female (i.e. AuDMV, Cg, CC, DLIVEnt, FrA, HIP, LO, MO, PtPR, PtA, PrL, M1, S1, RSC, M2, S2, TeA, TH, VO, and V1V2) mice that showed uncoupling at *p* < 0.05 levels (**Fig. 5D**). Importantly, at this age, male mice primarily showed a Type 2 uncoupled response in 15/28 brain regions, while females showed a Type 1 (increased perfusion, decreased glucose uptake) in 20/28 brain regions, further supporting what is known about APOE biology and sex dependent changes. These data are consistent with clinical findings [50], which showed similar regional uncoupling, and co-localization with TAU deposits measured via ^18^F-THK5317 PET, and corresponded to AD stage and severity [50].

Given the changes observed with APOE isoforms at 12 months, we then sought to determine if this approach could uncover neurovascular dysregulation at an earlier age. To assess the degree of metabolic and vascular dysfunction, we performed a regional analysis of neurovascular coupling [13, 48, 49] in both sexes at 8 months, which appeared to be a key transition age for AD progression based upon **Figs. 1-3**. Consistent with our hypothesis, B6 at 8 months showed an elevation of glucose uptake without a significant alteration in brain perfusion for most brain regions. Along with these changes, several key brain regions showed Type 2 uncoupling, which included: Cg, M1, M2, PtA, PrL, PtPR, and RSC. By contrast, male B6 mice at 8 months did not show significant numbers of regions that were uncoupled, and instead showed regional coupling which concurrently increased/decreased in metabolism and blood flow (**Fig. 6A**).

**FIGURE 6:**
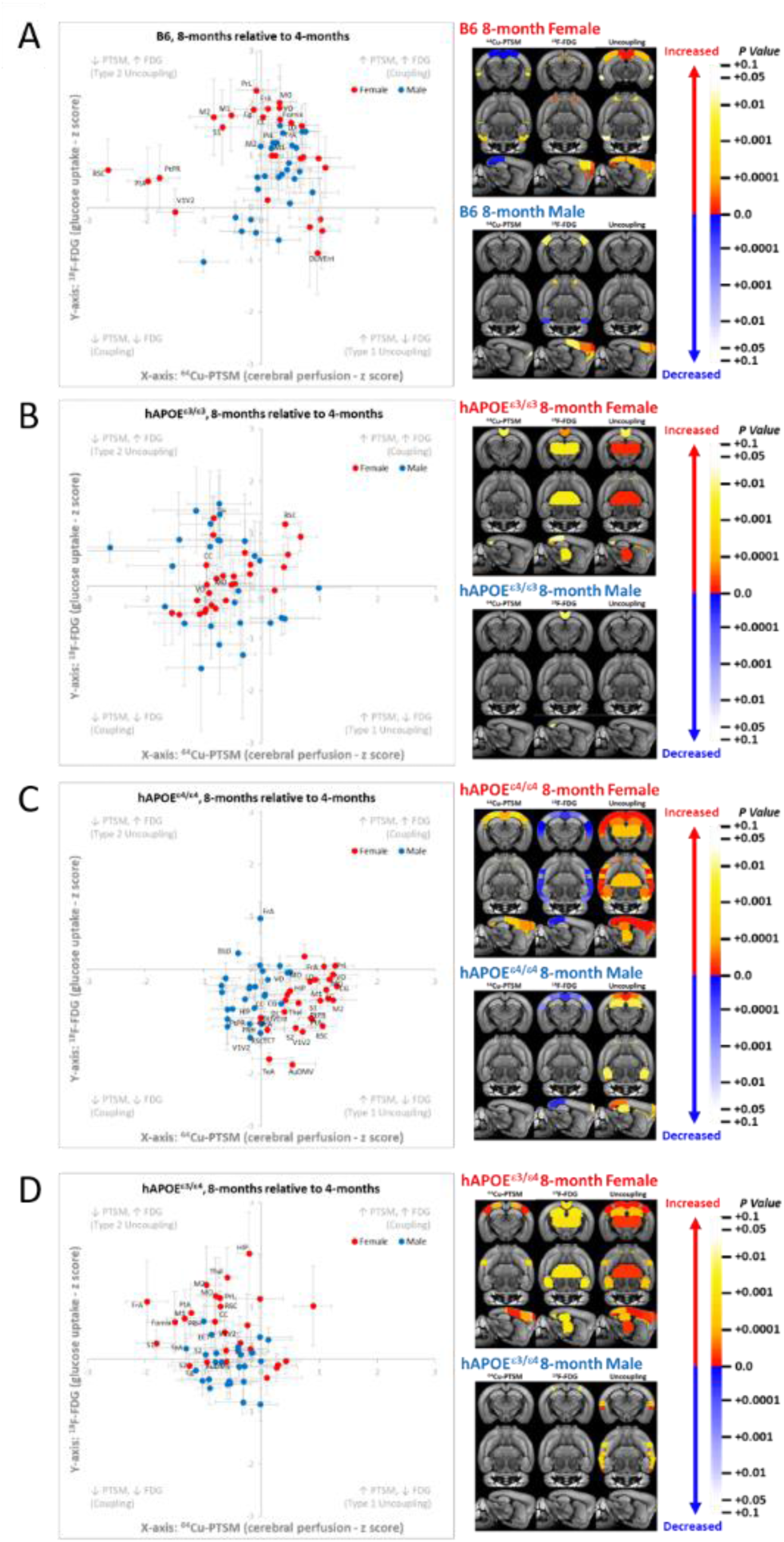
Assessment of Neurovascular Coupling at 8-months. (**A**) Analysis of neurovascular coupling of 8-month B6 in comparison with 4-month B6. Individual points represent the averaged z-scores of one of 27 brain regions, with red designating female and blue designating male. Image panels denote brain regions in the coronal, ventral and sagittal planes that exhibit significant differences in with each tracer, or the combined z-scores for uncoupling. Warmer colored regions indicated an increase, while cooler colored regions indicate a decrease. (**B**) Analysis of neurovascular coupling of 12-month h*APOE****^ε^****^3/^****^ε^****^3^* in comparison with 4-month h*APOE****^ε^****^3/^****^ε^****^3^*. (**C**) Analysis of neurovascular coupling of 12-month _h*APOE*_***ε****4/****ε****4* in comparison with 4-month h*APOE****^ε^****^4/^****^ε^****^4^*. (**D**) Analysis of neurovascular coupling of 12-month h*APOE****^ε^****^3/^****^ε^****^4^* in comparison with 4-month h*APOE****^ε^****^3/^****^ε^****^4^*.

Despite showing minimal changes at 12 months, we also examined neurovascular coordination in h*APOE****^ε^****^3/^****^ε^****^3^* in both sexes at the same 8 month time point. As with B6, the majority of the brain regions in both female and male mice showed a coupled phenotype, with the glucose uptake and perfusion showing coordinated increases/decreases. The few exceptions to this were CC, TH, MO, and VO in female mice which showed a Type 2 uncoupling of the perfusion and metabolism (**Fig. 6B**), which are white matter structures and are associated with synaptic integration and routing of sensory inputs.

Unlike the common isoform of APOE, the addition of h*APOE****^ε^****^4/^****^ε^****^4^* in 8 month mice demonstrated significant regional Type 1 uncoupling (i.e. metabolism decreasing and perfusion increasing) (**Fig. 6C**), with female mice showing both a larger number (i.e. 23/27 compared to 11/27) and greater distribution of uncoupled regions (i.e. AuDMV, CC, Cg, DLIVEnt, DI, ECT, FrA, HIP, LO, M1, M2,MO, PtPR, PtA, PRH, PrL, RSC, S1, S2, TeA, TH, VO, and V1V2) in circuits relevant to LOAD. By contrast, male mice at this age primarily showed a coupled phenotype with only 11/27 regions uncoupled (i.e. CC, Cg, DLO, FrA, HIP, MO, PtPR, PtA, RSC, VO, and V1V2).

Given the importance of allelic copy number and frequencies in the population, we were also able to conduct uncoupling analysis on h*APOE^ε3/ε4^* mice. We created these mice by intercrossing our h*APOE****^ε^****^3/^****^ε^****^3^* and h*APOE****^ε^****^4/^****^ε^****^4^*. At 8 months, much like the homozygous allelic counterpart, h*APOE^ε3/ε4^* showed a sexual dimorphism in the number and distribution of regions that showed uncoupling. However, specific to female h*APOE^ε3/ε4^* mice, the majority of key brain regions were involved in memory, learning, integration, and motor function (i.e. CC, Fornix, FrA, HIP, MO, PtA, PrL, M1, S1, RSC, M2, S2, TH, and V1V2) and exhibited a Type 2 uncoupled response (**Fig. 6D**). This elevated glycolytic state combined with a reduction in perfusion, coupled with transcriptomic (**Fig. 4**) and immunopathological (**Fig. 7**) changes in astroglial number and vascular density suggest that these brain regions may be uniquely more susceptible to cellular dysfunction.

**FIGURE 7:**
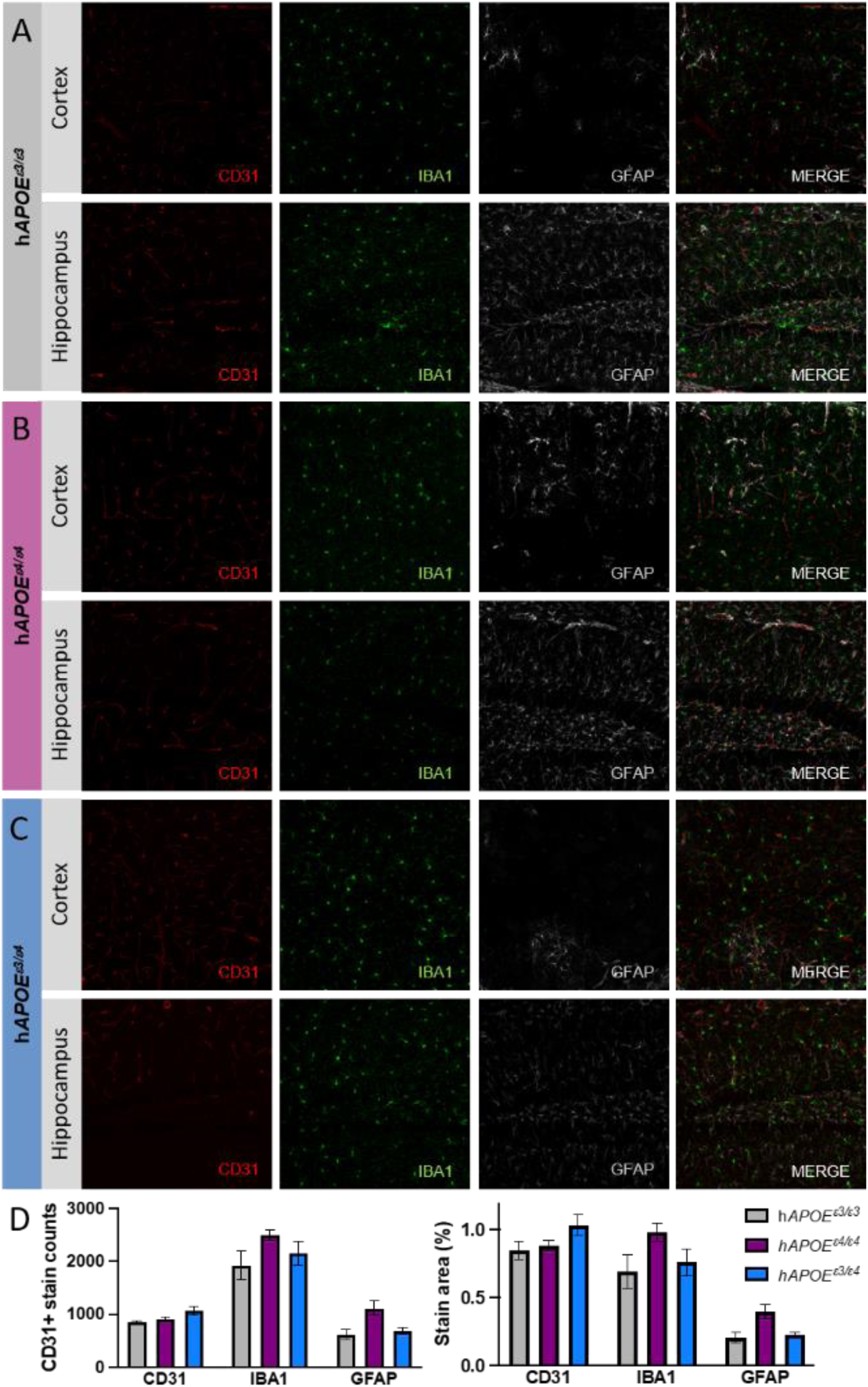
Neuropathology visualizing blood vessels, microglia and astrocytes. Panels (**A-C**) show representative images of CD31 (vessel density), IBA1 (activated microglia), and GFAP (activated astrocytes) in cortex (upper row) and hippocampus (lower row) across 8-month female h*APOE****^ε^****^3/^****^ε^****^3^* (grey; n=6, N=2), h*APOE****^ε^****^4/^****^ε^****^4^* (magenta; n=9, N=3) and h*APOE****^ε^****^3/^****^ε^****^4^* (blue; n=15, N=5) mice. Panel (**D**) show representative image quantitation of CD31, IBA1, and GFAP for all genotypes as cell counts (left) and percent area coverage (right).

### Neuropathology

One process outside of neural activity that has been suggested to contribute to differences in ^18^F-FDG signal, especially to increases, is neuroinflammation. A key predicted contributor to differences in ^64^Cu-PTSM signal is brain vascular density and integrity. As such, we sought to identify if there were significant differences across the different mouse h*APOE* strains imaged in this study by staining for vascular platelet endothelial cell adhesion molecule 1 (PECAM1) expression, microglia, astrocytes via CD31, IBA1, and GFAP, respectively (**Fig. 7A-C**). We focused on the 8-month time point in female as they showed the strongest phenotype from all genotypes included in our uncoupling analysis. Due to limited animal numbers, the power of these analysis was too low to detect changes between genotypes (**Fig. 7D**); however, trends were present which support our hypothesized cell specific contribution to Type 1 and Type 2 uncoupling. These data are consistent with previous reports for IBA1 [51, 52] and CD31 [53, 54].

## DISCUSSION

Our study was focused on using clinically relevant, translational measures to identify changes in glucose metabolism and cerebral perfusion in mice carrying humanized APOE variants across earlier stages of aging (8 and 12 months). This study did not include amyloid and tau as these models often have severe and accelerated neuropathology, limiting our ability to detect transitionary changes. We demonstrated regional differences in ^18^F-FDG or ^64^Cu-PTSM that were APOE isoform-, age- and sex dependent. We also combined ^18^F-FDG or ^64^Cu-PTSM PET measures (**Fig. 1-3**) with transcriptional profiling (**Fig. 4-6**), finding significant correlations between average signal of each tracer with differential gene expression and ontology. This study highlights the value of going beyond individual tracer analysis and standard general linear modeling (GLM) statistics, combining ^18^F-FDG and ^64^Cu-PTSM measures to assess neurovascular uncoupling across all 27 brain regions relative to age or a comparison group. Overall, we predict that this representation of imaging data will improve detection of early brain dysfunction and contribute to our ability to identify and expand the window in which preventive medicines can be effective.

At the start of this work, we were careful to include B6 mice for all studies, as this is the genetic background strain for the *APOE* allelic series. Numerous studies have highlighted the importance of genetic background for disease relevant phenotypes [55–57]. Until recently, much of the work examining h*APOE****^ε^****^4/^****^ε^****^4^* mice performed all comparisons to WT B6, which contain murine Apoe. Based on our work [56, 57], and the work of others [58], we determined that there are also significant differences between B6 and h*APOE****^ε^****^3/^****^ε^****^3^* mice. Some studies have suggested that mouse *Apoe* is more similar to h*APOE****^ε^****^4/^****^ε^****^4^* than h*APOE****^ε^****^3/^****^ε^****^3^*. In support of this, we found more brain regions that demonstrated significant differences in ^18^F-FDG and ^64^Cu-PTSM signal with age in B6 versus h*APOE****^ε^****^3/^****^ε^****^3^*, though the degree of uncoupled brain region metrics was relatively similar (**Fig. 1-2**, **5B**, **6A-B**). Our data suggest that mouse Apoe is functionally lies between h*APOE****^ε^****^3/^****^ε^****^3^* and h*APOE****^ε^****^4/^****^ε^****^4^* as it relates to neurovascular coupling.

Considering this, we then compared h*APOE****^ε^****^3/^****^ε^****^3^* and h*APOE****^ε^****^4/^****^ε^****^4^* mice across aging. While we also noted a sex effect in B6, this was even more apparent in h*APOE****^ε^****^3/^****^ε^****^3^*, and especially in h*APOE****^ε^****^4/^****^ε^****^4^* mice. Overall, female animals demonstrated the greatest number of regional changes in ^18^F-FDG and ^64^Cu-PTSM, as well as more regions that displayed neurovascular uncoupling (**Fig. 5C-D**, **6B-C**). In particular, female h*APOE****^ε^****^4/^****^ε^****^4^* animals exhibited severe decreases in ^18^F-FDG in brain regions such as the Cg, CC and RS, which are critical in the orchestration of widespread communication. The potential compensatory effect of blood flow through Type 1 uncoupling was highlighted with significant elevations in ^64^Cu-PTSM at 8 months, and this elevation was persistent through 12 months. However, the level of ‘maximum’ perfusion is also likely impacted by lack of change in vascular density observed with CD31 staining (**Fig. 7D**).

Based on our previous work [44], we included h*APOE****^ε^****^3/^****^ε^****^4^* mice in our assessment of neurovascular coupling at 8 months. Clinical studies often collapse *APOE****^ε^****^3/^****^ε^****^4^* and *APOE****^ε^****^4/^****^ε^****^4^* carriers together, and in the case of mouse studies, breeding restrictions on previous *APOE* allelic series made it challenging to generate h*APOE****^ε^****^3/^****^ε^****^4^* mice. However, the MODEL-AD *APOE* allelic series does not have the same breeding restrictions, allowing generation of h*APOE****^ε^****^3/^****^ε^****^4^* mice. Our data suggest that the underlying *APOE****^ε^****^4^* -dependent processes that increase risk for AD are not simply additive with allele copy, further supporting our previous study [44]. Here, we demonstrated that h*APOE****^ε^****^3/^****^ε^****^4^* females exhibit Type 2 uncoupling in multiple brain regions, a more severe and advanced disease phenotype than the Type 1 uncoupling observed in h*APOE****^ε^****^4/^****^ε^****^4^* mice at 8 and 12 months. Furthermore, while ^18^F-FDG uptake appeared to be more heavily impacted with age in most strains, ^64^Cu-PTSM was the predominant measure that changed in h*APOE****^ε^****^3/^****^ε^****^4^*. These data are supported in h*APOE****^ε^****^3/^****^ε^****^4^*, which have the highest amount of CD31 staining relative to h*APOE****^ε^****^3/^****^ε^****^3^* mice (**Fig. 7D**). This more severe uncoupling at a key transition age may point to differences in overall compensatory networks and cells involved [16, 59]. For instance, if ones genetic make-up is h*APOE****^ε^****^3/^****^ε^****^3^* or h*APOE****^ε^****^4/^****^ε^****^4^* from birth, this may result in immunological adaptation throughout development, as has been observed with the complement system [60]. By contrast, mice, which carry h*APOE****^ε^****^3/^****^ε^****^4^*, may develop a negative allelic interaction, which potentiate the immune response [61]. Importantly, our previous work identified DE that encode for hormone and insulin signaling in mice that carry h*APOE****^ε^****^3/^****^ε^****^4^* [44], and are consistent with the Type 2 uncoupling seen in the current work.

To our knowledge, our study was the first to correlate PET tracer and gene expression changes. While this analysis had to be performed on a brain-wide scale due to averaging of z-scores across all 27 brain regions, there were still numerous DE that showed significant positive and negative correlations. We were also able to perform GO term analysis on DE, which highlighted a number of pathways directly relevant to cerebral metabolism and perfusion. For example, significant correlations with ^18^F-FDG across groups highlighted metabolic and neuronal pathways, and are consistent with previous work, which demonstrated metabolic reprogramming of cellular metabolism with age and genotype [9, 62, 63]. In addition, significant correlations with ^64^Cu-PTSM netted pathways relevant to ‘tight junction assembly’ and ‘cell migration’. For comparisons across age in h*APOE****^ε^****^4/^****^ε^****^4^*, ‘ADP metabolic processes appeared as a significant GO term. While on its face ^18^F-FDG may appear to be a more direct correlate, ADP represents a critical metabolic intermediate [10, 11], and purinergic signaling molecule [64–67], stored inside of blood platelets. Moreover, ADP is directly phosphorylated to synthesize ATP via glycolysis and oxidative phosphorylation in the cell [10, 11], or can be converted directly to adenosine via CD39 and CD73 [68], resulting in blood vessel dilation and increases in tissue perfusion [69].

The overall focus of our work was to understand differences in cerebral metabolism and perfusion across aging and *APOE* allelic combinations using translational measurements. While we did include traditional analyses in this study, we were not able to directly detect mechanisms underlying these differences using standard GLM statistical modeling approaches. However, the transcriptomic (**Fig. 4**), uncoupling (**Figs. 5-6),** immunopathology (**Fig. 7**) findings presented here in combination with the supporting literature enables us to speculate the underlying mechanisms driving Type 1 and Type 2 neurovascular uncoupling, and predict how this may be coordinated to stage of disease. For instance, Type 1 neurovascular uncoupling is thought to be driven by a cytokine (i.e. TNFα, IL1β, IL6 and/or IL12) [70–73] down-regulation of insulin receptors [74, 75], which in turn results in an apparent reduction in neuronal glucose uptake via GLUT3 [63, 76, 77], ensuing a metabolic deficit (**Fig. 8**). This deficit is partially offset by astrocytic uptake of glucose via GLUT1 [76, 77], which is converted to lactate and shuttled to neurons (i.e. lactate shuttle) [77, 78] via MCT2 export on astrocytes and subsequent uptake via MCT1 and 4 [76, 77].

**FIGURE 8:**
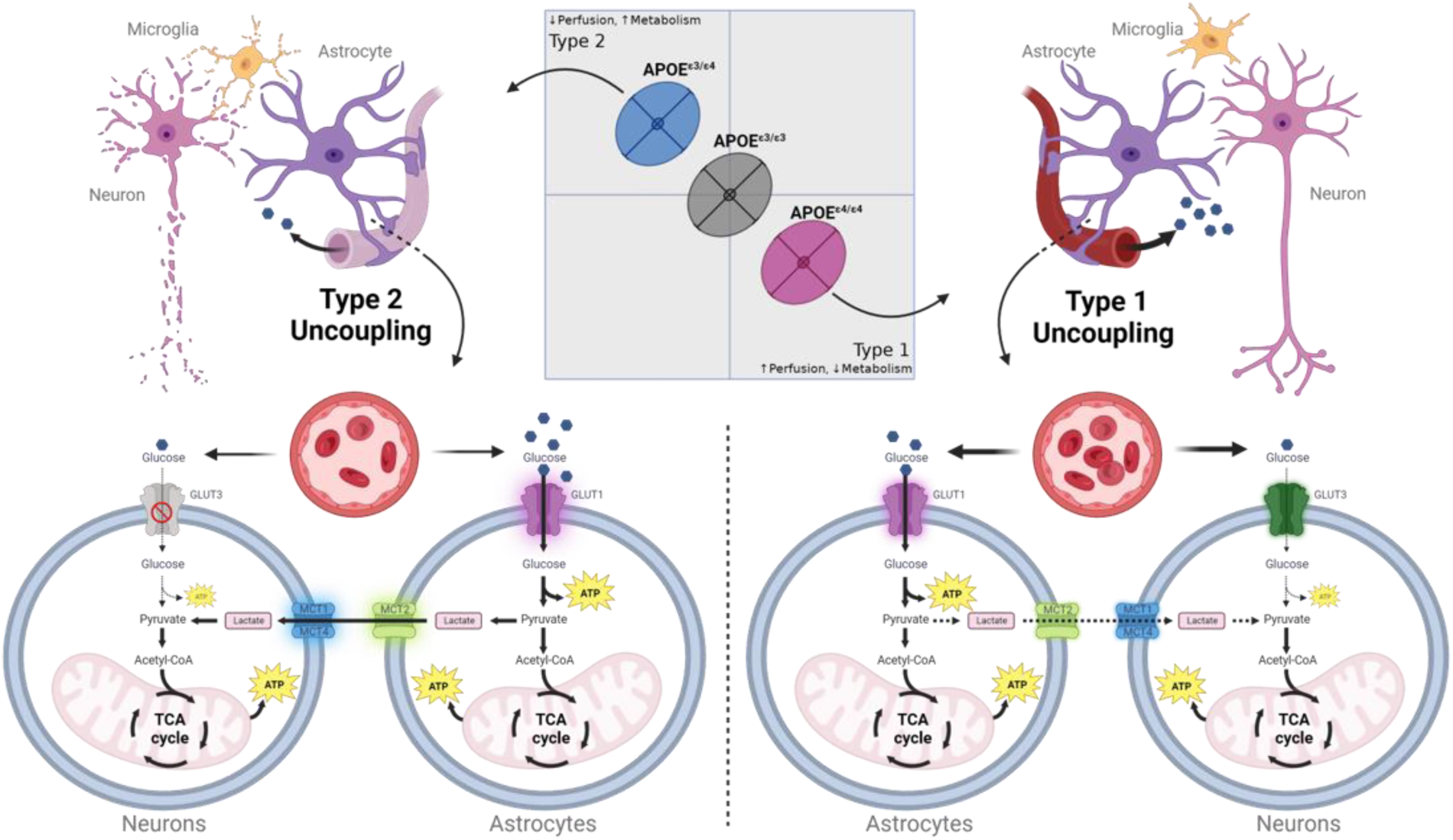
Predicted mechanisms underlying Type 1 and Type 2 Uncoupling in Alzheimer’s disease risk. (**A**) Graphic depicting Neurovascular Coupling analysis for *APOE* allelic series. The predicted underlying cell types, molecules and receptor response relevant to Type 1 and Type 2 uncoupling are depicted.

Commensurate with these changes, we hypothesize compensatory vasodilation of the vascular smooth muscles, which results in an apparent increase in tissue perfusion, or reactive hyperemia (**Figs. 4-7**). Furthermore, we hypothesize that sustained elevations in cytokine release (i.e. TNFα, IL1β, IL6, and/or IL12) [70–73] results in Type 2 Neurovascular Uncoupling, leading to a cessation of GLUT3 transport on neurons, and a cytokine driven upregulation of GLUT1 receptors on astrocytes (**Fig. 8**) and resulting astrogliosis [79]. This net increase in transport capacity, we hypothesize, results in greater astrocytic glucose uptake to support neuronal metabolism via the lactate shuttle via MCT2 and MCT1 and 4 transport [76, 77]. Additionally, we hypothesize that sustained cytokine release will result in astroglial damage of the vascular unit, resulting an inability of the vasculature to increase perfusion further and blood brain barrier dysfunction [80–82]. Overall, this results in a mismatch between supply and demand observed in our work (**Figs. 4-7**).

As with all research, there are limitations, which should be considered when developing new analytical approaches. The current study employs a variant on the z-score analysis for a single factor that extends this in two dimensions, and then projects this onto a Cartesian coordinate system for ease of visualization and categorization. Because of this, all uncoupling plots represent relative changes to a reference population (i.e. age or genotype), and therefore it is critical that this reference group be matched for tracer, sex, and the factor of interest before plotting (see **Figs. 5-6**). In addition, although this approach has been applied in this context to PET data, it should be noted that because the z-score transformations remove units and scale, this approach could be applied in any multi-modal context (PET, CT, or MRI) to assess dysregulation of perfusion and metabolism.

Finally, future work will explore the underlying mechanisms, which are hypothesized above, to determine how and which cell types, transporters, and metabolites play a role in these measures. In addition, because of the universality of this approach, future work would benefit from aligning mouse to human studies as a means to identify which brain regions are most susceptible to dysregulation and may progress to a more advanced disease stage.

## DATA AVAILABILITY STATEMENT

The contributions presented in the study are publicly available. The imaging data are available via the AD Knowledge Portal: https://adknowledgeportal.org (permission was obtained for this material through a Creative Commons CC-BY license). Data are available for general research use according to the following requirements for data access and data attribution (https://adknowledgeportal.org/DataAccess/Instructions).

## AUTHOR CONTRIBUTIONS

KO, PBL, RSP, SAP, CPB, EWM, KE, JNK, KEF, GWC, GRH, and PRT contributed to conception, design, and performance of studies and data analysis. PBL, CPB, SAP, and PRT developed and tested the analysis methods and templates. KO, RSP, KEF, GRH, and PRT wrote the first draft of the manuscript. KO, CPB, SAP, and PRT performed the statistical analysis. All authors contributed to the manuscript revisions, read, and approved the submitted version.

## CONSENT STATMENT

No human subjects were used in the current study, and therefore consent was not necessary.

## ACKNOWLEDGEMENTS

The authors would like to thank Kelly Keezer for generating animals at The Jackson Laboratory and shipping them to Indiana University. We would also like to thank Amanda Hewes for imaging immunopathology brain sections. In addition, Adrian Oblak for providing animals and Amanda Bedwell for project organization at Indiana University. We would also like to recognize, The Jackson Laboratory Genome Technology Core, and The Jackson Laboratory Microscopy Core for their contributions to this study.

