## Supplemental Materials for "Assessment of Neurovascular Uncoupling: *APOE* Status is a Key Driver of Early Metabolic and Vascular Dysfunction"

| Index | Region of Interest (ROI) Name | ROI Label |
|---|---|---|
| 1 | Agranular Insular Cortex (Doral-Ventral) | AI |
| 2 | Auditory Cortex (Dorsal, Medial, Ventral) | AuDMV |
| 3 | Caudate Putamen (Dorsal Striatum) | CPu |
| 4 | Cingulate Cortex | Cg |
| 5 | Corpus Callosum | CC |
| 6 | Dorosolateral Orbital Cortex | DLO |
| 7 | Dorsintermed Entorhinal Cortex | DLIVEnt |
| 8 | Dysgranular Insular Cortex | DI |
| 9 | Entorhinal Cortex | ECT |
| 10 | Fornix | Fornix |
| 11 | Frontal Association Cortex | FrA |
| 12 | Hippocampus (CA1-CA3) | HIP |
| 13 | Lateral Orbital Cortex | LO |
| 14 | Medial Orbital Cortex | MO |
| 15 | Parietal Corext (Post-Rostral) | PtPR |
| 16 | Perietal Association Cortex (Lateral-Medial) | PtA |
| 17 | Perirhinal Cortex | PRH |
| 18 | Prelimbic Cortex | PrL |
| 19 | Primary Motor Cortex | M1 |
| 20 | Primary Somatosenory Cortex | S1 |
| 21 | Retrosplenial Dysgranular Cortex | RSC |
| 22 | Secondary Motor Cortex | M2 |
| 23 | Secondary Somatosenory Cortex | S2 |
| 24 | Temporal Association Cortex | TeA |
| 25 | Thalmus | TH |
| 26 | Ventral Orbital Cortex | VO |
| 27 | Visual Cortex (Primary and Secondary) | V1V2 |

*Supplementary Table 1: Full names of network nodes (region of interest) as defined in the Paxinos and Franklin Atlas. Left and right regions were averaged to yield a network of 27 regions normalized by the cerebellum.*
